# Intraluminal neutrophils limit epithelium damage by reducing pathogen assault on intestinal epithelial cells during *Salmonella* gut infection

**DOI:** 10.1101/2023.02.23.529832

**Authors:** Ersin Gül, Stefan A. Fattinger, Bidong D. Nguyen, Annika Hausmann, Markus Furter, Manja Barthel, Mikael E. Sellin, Wolf-Dietrich Hardt

**Affiliations:** Institute of Microbiology, Department of Biology, ETH Zurich, Zurich, Switzerland; Science for Life Laboratory, Department of Medical Biochemistry and Microbiology, Uppsala University, Uppsala, Sweden

**Author notes:** **<u>Correspondence:</u>**.

## Abstract

Recruitment of neutrophils into the gut epithelium is a cardinal feature of intestinal inflammation in response to enteric infections. Previous work using the model pathogen *Salmonella* Typhimurium (*S*. Tm) established that invasion of intestinal epithelial cells by *S*.Tm leads to recruitment of neutrophils into the gut lumen, where they can reduce pathogen loads transiently. Notably, a fraction of the pathogen population can survive this defense, re-grow to high density, and continue triggering enteropathy. However, the functions of intraluminal neutrophils in the defense against enteric pathogens and their effects on preventing or aggravating epithelial damage are still not fully understood. Here, we address this question via neutrophil depletion in different mouse models of *Salmonella* colitis, which differ in their degree of enteropathy. In an antibiotic pre-treated mouse model, neutrophil depletion by an anti-Ly6G antibody exacerbated epithelial damage. This could be linked to compromised neutrophil-mediated elimination and reduced physical blocking of the gut-luminal *S*.Tm population such that the pathogen density remained high near the epithelial surface throughout the infection. The removal of luminal *S*. Tm by gentamicin, an antibiotic restricted to the gut lumen, reversed the effect of neutrophil depletion on epithelial cell loss. Strikingly, when using germ-free mice and an *S*. Tm *ssaV* mutant capable of epithelium invasion, but attenuated for survival and growth within host tissues, neutrophil depletion caused exacerbated immune activation of the gut mucosa and a complete destruction of the epithelial barrier. Together, our data indicate that intraluminal neutrophils are central for maintaining epithelial barrier integrity during acute *Salmonella*-induced gut inflammation, by limiting the sustained pathogen assault on the epithelium in a critical window of the infection.

**Highlights:**

○ After the first wave of mucosal invasion (day 1 p.i.), *S*. Tm maintains the assault from the lumen, triggering the continued expulsion of epithelial cells in antibiotic pre-treated mice.
○ Neutrophil recruitment into the gut lumen is essential to limit this continued *Salmonella* attack on the epithelium.
○ In antibiotic pre-treated SPF mice, neutrophil depletion exacerbates *S*. Tm invasion, causing excessive epithelial cell loss, which compromises epithelial barrier integrity at later time points (day 2-3 p.i.).
○ In germ-free mice, neutrophil depletion exacerbates epithelial responses and epithelial barrier destruction even more strongly than in streptomycin pre-treated SPF mice.
○ Gentamicin treatment and *ssaV* mutant infections indicate that neutrophils prevent epithelial damage by eliminating and physically blocking gut-luminal pathogens.

## Introduction

Polymorphonuclear leukocytes (PMN), also called neutrophils, are the most abundant immune cell type in circulation and considered as an early line of defense against many infections, including those caused by enteropathogenic bacteria. Neutrophil recruitment into and across the gut epithelium is a hallmark of infectious (e.g., pathogen-induced) and non-infectious (e.g., inflammatory bowel disease) colitis [1-6]. They provide a powerful defense during enteric infections using a myriad of effector mechanisms (i.e., reactive oxygen/nitrate species (ROS/RNS), antimicrobial peptides (MPO/NE), and neutrophil extracellular traps (NETs) [7, 8]. While being indispensable in the defense against microbes, neutrophils are also often associated with tissue damage during gut inflammation [5]. Therefore, the degree of protective versus tissue damaging effects of neutrophils is highly context dependent and is not fully understood for all natural infections and infection models traditionally used in the field.

Intestinal inflammation caused by enteric pathogens such as Non-typhoidal *Salmonella enterica* serovars (NTS), including serovar Typhimurium (*S*.Tm) has been extensively studied in recent years. Murine models of *S*.Tm gut inflammation such as the Streptomycin pre-treatment model provide an excellent basis to study the stages of gut inflammation during *Salmonella* infection [9, 10]. To elicit disease in streptomycin pre-treated mice, *S*.Tm invades intestinal epithelial cells using its type-three secretion systems (TTSS)-1 and -2 [2]. Invasion of the epithelial cells results in activation of the NLRC4 inflammasome. This results in two major outcomes: i) expulsion of infected epithelial cells into the gut lumen and ii) recruitment of innate immune cells including neutrophils via secretion of IL-18 and other immune mediators (Müller et al., 2016; Sellin et al., 2014). Studies in this model suggest that neutrophils may have stage-specific functions during the acute infection. In the first 18h after an oral dose of *S*.Tm, neutrophils appear not to be involved in defense against *Salmonella* [11]. However, follow-up research using the same model showed that Gr-1+ cells (neutrophils and monocytes) impose a strong bottleneck, reducing the gut luminal *S*.Tm population to as little as 15’000 *S*.Tm cells by 48-72h p.i. [12]). Interestingly, the pathogen population can grow back by 96h p.i. (to approx. 10^9^ CFU / g feces). However, it remains unclear if these high pathogen densities in the gut lumen may represent an important continued driver of enteropathy at that stage of the infection as well. Similarly, due to their stage-specific effects, it remains unclear whether the recruitment of neutrophils is beneficial or rather a detriment for gut mucosal integrity during acute *Salmonella* infection.

Infected epithelial cells are expelled during enteric diseases to prevent the subsequent transmigration of the pathogens into the underlying tissue (lamina propria; LP) and the systemic organs [13-17]. Moreover, the expulsion may limit the amounts of inflammasome-dependent cytokines like IL-18 which are released into the lamina propria. The resulting immune responses lead to temporary shortening of the intestinal crypts and require the rapid division of crypt stem cells to retain barrier integrity (crypt hyperplasia; [10, 15, 18]). A failure to replace the expelled epithelial cells, as reported in germ-free mice infected with wild type *S*. Tm [18], can lead to shortened crypt structures and can be detrimental to epithelial barrier integrity. Similarly, excessive assault on epithelial cells and increased pathogen tissue loads due to the lack of one or more of the mucosal defenses can also drive pathological cell loss and destruction of the epithelium. This was shown in the examples of NAIP/NLRC4 or GSDMD (downstream of NLRC4 inflammasome; executer of pyroptotic cell death [19]) deficiency, where failure to initiate timely epithelial immune responses resulted in increased pathogen loads and disruption of the epithelium at later stages of the infection [20, 21]. These reports highlight the importance of a balanced and timely mucosal immune response to maintain epithelium integrity during enteric infections. However, as wild type *S*. Tm colonizes the gut lumen, the gut epithelium and the lamina propria, it remained unclear which of these pathogen populations would be targeted by the neutrophil defense and how this affects the epithelial barrier. We hypothesized that neutrophils might provide an additional layer of defense in the gut lumen and alleviating the burden on the gut epithelium during severe stages of acute *Salmonella* infection.

Here we used a comprehensive approach to assess the function of intraluminal neutrophils during *Salmonella*-induced gut inflammation. Utilization of mouse models with varying severity of *S*.Tm-induced colitis allowed us to highlight the stages of gut inflammation where neutrophils exert a crucial epithelium-protective function in the infected gut.

## Results

### Neutrophils slow down the progression of cecal tissue infection by day 3 of *S*.Tm infection

We previously showed that Gr-1+ cells (neutrophils and monocytes) contribute to the control of the gut-luminal pathogen population in the streptomycin mouse model for acute *Salmonella* gut infection. This defence eliminates ≈99.999% of the gut luminal *S*.Tm population by day 2 p.i. [12]. Interestingly, the luminal pathogen population can grow back up to ≈10^9^ CFU / g stool by days 3-4 p.i. However, it remained unclear whether the tissue-lodged or the re-growing luminal pathogen population contribute to the enteropathy at this “mature” stage of the infection. Similarly, the role of intraluminal vs tissue-lodged neutrophils during this stage of *Salmonella* gut infection was still an enigma. Here, we addressed these topics and specifically focused on the consequences of neutrophil depletion on the epithelial barrier, which has previously been shown to respond to *S*.Tm onslaught by a sensitive epithelial cell expulsion response [13, 15, 21, 22]. To explore this, we orally infected streptomycin pretreated C57BL/6 mice with 5×10^7^ CFU of wild-type *S*.Tm (SL1344) for 3 days. To detect possible bottlenecks on the luminal pathogen population, we included seven wild-type isogenic tagged *S*. Tm strains in the inoculum (WITS; [23, 24]) at a ratio of 1:1000 (tagged: untagged). This ratio of tagged versus untagged wild-type strains will result in the random loss of abundance in one or more of the tagged strains if the gut-luminal *S*.Tm population undergoes a transient bottleneck. Importantly, the resulting unequal tag-distribution will be maintained within the luminal *S*.Tm population, even after the population has re-grown to carrying capacity at day 3 p.i.. Therefore, qPCR analysis of the tag abundances allows us to quantitatively assess gut luminal bottlenecks in the streptomycin pre-treatment mouse model [12]. To assess the role of neutrophils specifically, we treated mice with a neutrophil-depleting antibody directed against Ly6G (α-Ly6G; cl. 1A8; via intraperitoneal injection (I.P.); see details in **Materials & Methods**). This method reduces the number of neutrophils recruited to the gut tissue during *Salmonella* infection (**Fig. 1A**; [11]). In control experiments, we infected the mice with the same protocol but treated them with PBS only. In line with our previous studies, neutrophil depletion did not change the pathogen densities in the feces or systemic nor did it cause a bottleneck on the gut luminal *S*.Tm population (both groups; evenness score: ∼1) during the first stage of the acute disease (first 2 days p.i.; **Fig. S1A-D**; [12]). At a later stage (day 3 p.i.), on the contrary, 24 out of 77 WITS were lost in the control (evenness score: ∼0.46; **Fig. 1B-C**), while neutrophil-depleted mice retained a much higher fraction of all tags (only 3 out of 49 WITS were lost with an evenness score: ∼0.92; **Fig. 1B-C**), confirming the role of neutrophils in this previously observed bottleneck effect. This was similar to our previous observations with the Gr-1-based depletion (anti-Gr-1 antibody; both monocytes and neutrophils depleted). Furthermore, the luminal pathogen densities in the cecal content were significantly higher in mice with neutrophil depletion compared to the control mice both at day 2 and at 3 p.i. (**Fig. S1E**) as we also showed before [12]. In conclusion, these data reproduced the role of neutrophils in inflicting a transient bottleneck on the gut luminal *S*. Tm population and provided us with materials to assess the effects of the neutrophil depletion on the enteropathy, which had not been assessed in the earlier work.

**Fig. 1.**
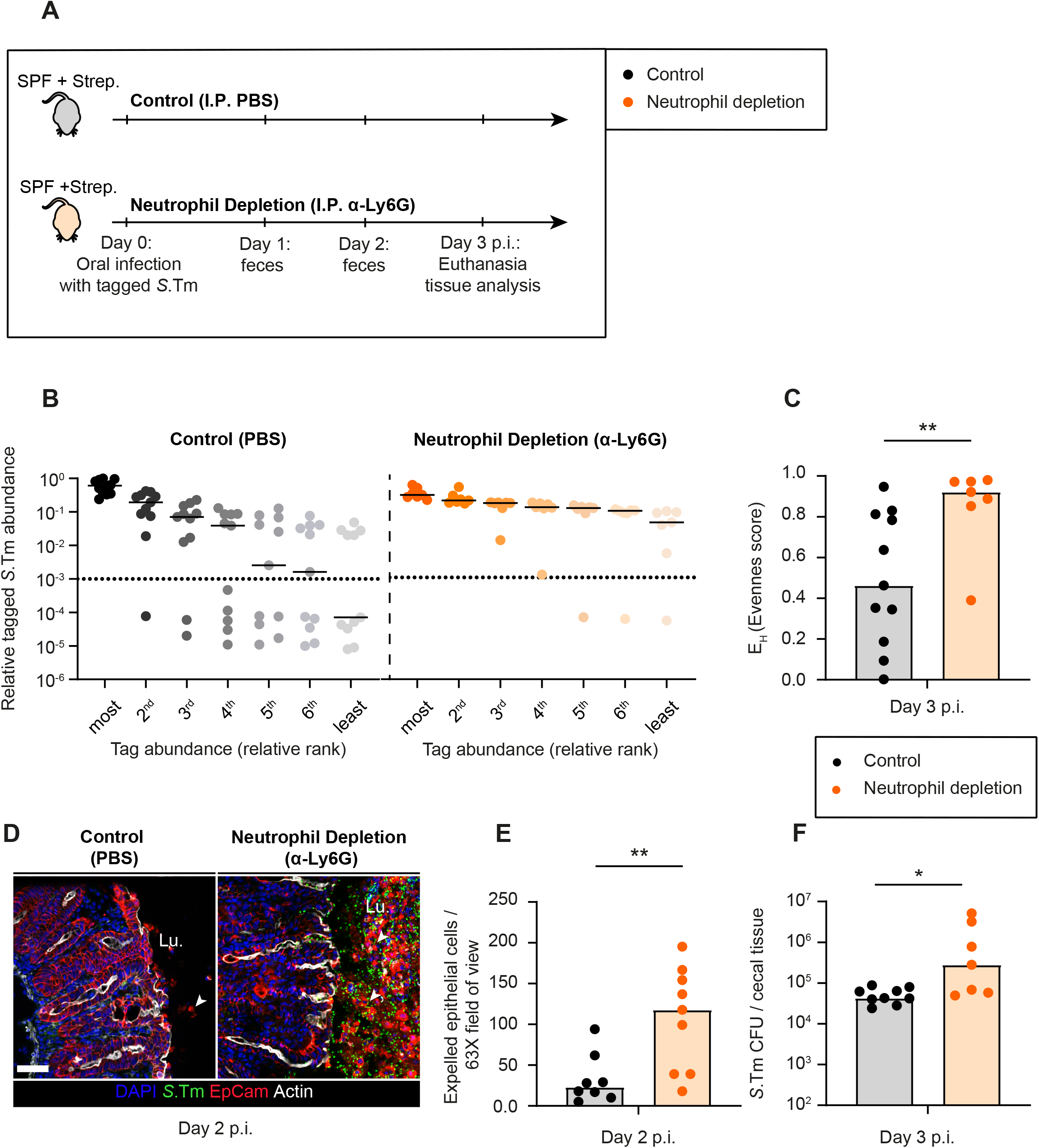
Consequences of neutrophil depletion with α -Ly6G on luminal pathogen loads and epithelial cell expulsion. **A)** Scheme summarizing the experimental setup of Panels **B-F**. Streptomycin pre-treated C57BL/6 mice were infected orally with 5×10^7^ CFU of wt *S*.Tm (SL1344) for 3 days. One group (control) treated with the vector (PBS; black symbols) and the second group with α-Ly6G (orange symbols) intraperitoneally (I.P.). **B)** Relative ranks of the tagged *S*.Tm strain abundances in feces at day 3 p.i.. **C)** Evenness score at day 3 p.i. **D)** Representative micrographs of cecal tissue sections, stained for epithelial marker EpCam and *Salmonella* LPS. Lu. = Lumen. White arrows point at expelled epithelial cells. Scale bar=50 µm. **E**) Microscopy-based quantification of IECs per 63x field of view (i.e., cells/high power field; hpf). Each data point is the average of 5 fields of view (FOV) per section. **F)** *S*.Tm pathogen loads in cecal tissue (CFU / g). Lines or Upper ends of the bars indicate the median. Dotted lines indicate the detection limit. **Panels B-C**) Pooled from 2 independent experiments for each group: control (n=11 mice) and neutrophil depletion (n=7 mice). **Panels D-F**) Pooled from 3 independent experiments for each group: control (n=8 mice) and neutrophil depletion (n=9 mice). Two-tailed Mann Whitney-U tests were used to compare two groups in each panel. p≥0.05 not significant (ns), p<0.05 (*), p<0.01 (**).

To assess the degree of enteropathy at later stages, we focused on the cecal tissue (main site of invasion in streptomycin pretreated model [10]). Strikingly, fluorescence microscopy of cecal tissue at day 2 p.i. revealed massive epithelial shedding in mice with neutrophil depletion, while the control mice showed only a few epithelial cell expulsion events. (**Fig. 1D-E**). The mice with neutrophil depletion had on average 5-fold more expelled cells in the cecum lumen and showed disrupted crypt structure, whereas the control mice featured the typical infection-associated crypt hyperplasia. (**Fig. 1D-E**). Of note, at this stage, epithelial cells often dislodged from the mucosa in the form of large cell aggregates, which is different from the typical NAIP/NLRC4-driven epithelial cell expulsion that we and others described before, where single infected cells are selectively extruded by their neighbours [13, 21, 22]. However, the underlying causes remained unclear.

We reasoned that in the absence of neutrophils, epithelial cells might be subject to more *S*.Tm invasion events as the gut luminal pathogen loads remained higher than in the non-depleted controls during the bottleneck-phase of the infection (i.e. day 2 p.i.). This might result in higher net pathogen invasion into the gut tissue. To test this, we quantified the pathogen loads in the cecal tissue using the gentamicin protection assay. Indeed, the number of *S*.Tm per cecal tissue was elevated in neutrophil-depleted mice in comparison to the control mice at day 3 p.i. (**Fig. 1F**). The elevated tissue invasion frequency was also reflected in spleen and liver (**Fig. S1F**). However, pathogen loads in the mLN did not differ significantly between the depleted mice and the controls (**Fig. S1F**). The reasons for this remain unclear, but this observation could be linked to the balance between pathogen migration into the mLN, the pathogen growth in that organ and the antimicrobial defense mounted dynamically to control pathogen growth at that site [23]. Taken together, these observations suggest that in the absence of neutrophil defense, the gut mucosa experiences higher *S*.Tm loads at ∼2-3 days of infection, coupled to excessive and uncontrolled shedding of intestinal epithelial cells into the lumen.

### Neutrophils provide a physical barrier against *S*.Tm invading from the gut lumen

It was shown that following an acute *Toxoplasma gondii* gastrointestinal infection of mice, neutrophils can form intraluminal casts that contain commensal outgrowth and prevent spread of pathobionts to the systemic organs [25]. Besides, neutrophils have been shown to interact with luminal *S*.Tm closely during the early stages (at 20h p.i.) of our streptomycin mouse model [1]. Therefore, we speculated that neutrophils might not only be killing luminal pathogens, but also may form a physical barrier to reduce *S*.Tm access to the epithelial surface, thereby restricting further pathogen invasion events. Fluorescence microscopy of cecal tissue at day 1 and 3 p.i. revealed that intraluminal neutrophils and luminal *S*.Tm were in close contact in streptomycin pretreated mice (**Fig. 2A**). At day 1 p.i., neutrophils were present in the gut lumen in aggregates of various sizes (ca. 2 cells to 50 cells). We asked whether there was a correlation between the size of the neutrophil aggregates in the lumen and the number of *S*. Tm cells interacting with the epithelium at this time point (**Fig. 2B**). To quantify this, we calculated the median number of neutrophils per 63-X field of view and compared the number of *S*.Tm cells interacting with the epithelium in two scenarios, that is in fields of view with >15 neutrophils vs fields of view with ≤15 neutrophils. Strikingly, the number of *S*.Tm cells in close association with the epithelium was significantly higher in the scenario where there were only a few (≤15) neutrophils in the area in comparison to the scenario with >15 neutrophils (**Fig. 2B**). Of further note, we observed that the number of neutrophils in the gut lumen was significantly higher at day 3 p.i. compared to day 1 p.i. (**Fig. 2C**). At day 3 p.i., neutrophils assembled into dense structures in front of the epithelium reminiscent of the intraluminal casts reported in other contexts [25]. In accordance with this, the average numbers of epithelium-associated pathogen cells were significantly lower at day 3 p.i. compared to day 1 p.i. (**Fig. 2D**). Altogether, our findings suggest that in response to *S*.Tm infection neutrophils are recruited to the gut lumen where they both kill *S*.Tm, resulting in a population bottleneck, and at the same time generate a physical barrier against *Salmonella* cells building up from day 1 p.i. onwards in the streptomycin pre-treated mouse model.

**Fig. 2.**
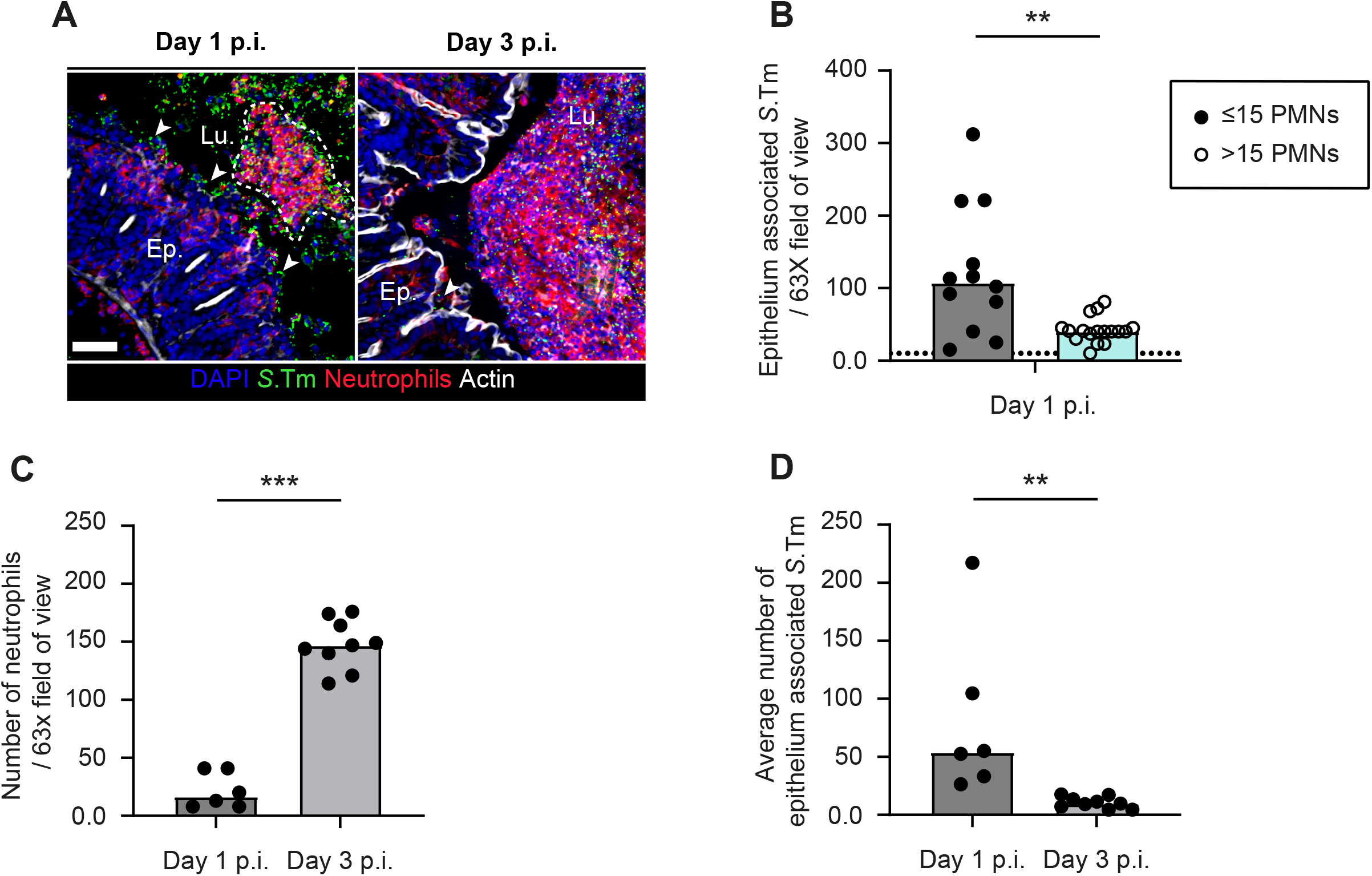
Microscopy analysis of the time course of neutrophil infiltration into the gut lumen (day 1 & 3 p.i.) **A)** Representative micrographs of cecal tissue sections from mice infected with *S*.Tm, taken at day 1 and 3 p.i., stained for neutrophil marker Ly6B.2 and *Salmonella* LPS. Lu.=Lumen. Ep.=Epithelium. White arrows indicate *S*.Tm associated with the epithelium. Scale bar=50 µm. **B-D)** Microscopy-based quantification of **B)** *S*.Tm associated with the epithelium at day 1 p.i.. (total 24 FOVs from 6 mice; filled symbols; neutrophils per field≤15 vs empty symbols; neutrophils per field>15). **C)** neutrophils per 63x field of view (each data point is the average of 5 FOVs per section of the same mouse). **D)** average number of *S*.Tm associated with the epithelium. Upper ends of the bars indicate the median. Dotted lines indicate the detection limit. **Panels C-D**) Pooled from 3 independent experiments for each group: day 1 p.i. (n= 6 mice) and day 3 p.i. (n=9 mice). Two-tailed Mann Whitney-U tests were used to compare two groups in each panel. p≥0.05 not significant (ns), p<0.01 (**), p<0.001 (***).

### Neutrophils release extracellular traps (NETs) into the gut lumen in response to acute *Salmonella* infection

Next, we investigated the nature of the neutrophil barrier(s) against pathogen attack. Neutrophils are equipped with an arsenal of anti-microbial effector mechanisms including phagocytosis, release of reactive oxygen/ nitrogen species (ROS/RNS), granules filled with antimicrobial agents, and can in addition release neutrophil extracellular traps (NETs; [7, 8, 26]). NETs are networks of extracellular host-DNA decorated with antimicrobial peptides. They are released in response to several stimuli and can kill or immobilize the pathogens in the extracellular space [27]. We speculated that the release of NETs into the gut lumen could provide one mechanism by which neutrophils limit further *S*.Tm invasion into the cecum epithelium. Since their discovery two decades ago [26] as a powerful and unorthodox immune defense mechanism, NETs have been implicated not only in the defense against many extracellular pathogens [27]), but also in tissue damage and autoinflammatory diseases [5, 28, 29]. We reasoned that NETs-entrapped *S*.Tm cells might not only be immobilized, but might also be efficiently removed with the fecal flow. NETs are generated during a programmed cell death known as ‘NETosis’, which is initiated by peptidyl arginine deiminase-4 (PAD4, aka Padi4), promoting the formation of citrullinated histone 3 (cit-His3). A previous study showed that PAD4 is necessary for an efficient neutrophil response to the enteric pathogen *Citrobacter rodentium* [30]. Therefore, cit-His3 can be used as a marker for NETs. Our microscopy analysis of intraluminal neutrophil aggregates at day 3 p.i. indeed revealed that parts of these aggregates contained DNA with citrullinated histones (**Fig. 3A**). Interestingly, these cit-His3+ regions showed an overlap with regions where *S*.Tm cells were densely localized in the lumen. These results suggest that intraluminal neutrophils can undergo NETosis during acute *Salmonella* infection to kill/immobilize the pathogen and reduce further attacks onto the epithelium.

**Fig. 3.**
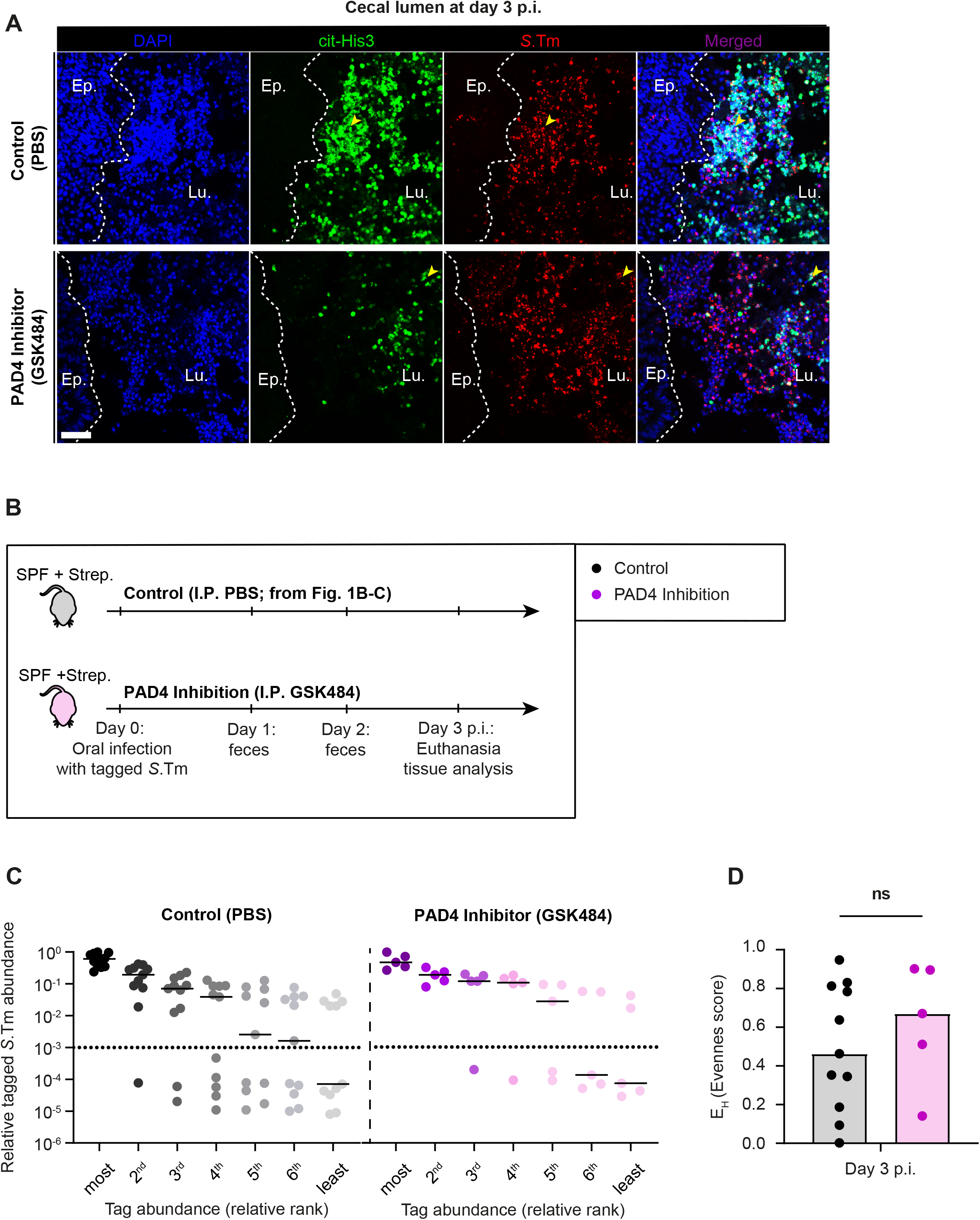
Microscopy analysis of intraluminal NETs at day 3 p.i. **A)** Representative micrographs of cecal tissue sections from mice infected with *S*.Tm, taken at 3 days p.i., stained for NET marker cit-His3 and *Salmonella* LPS. Lu. = Lumen. Ep. = Epithelium. Yellow arrows indicate *S*.Tm associated with the NET marker cit-His3. White dotted lines indicate the epithelial barrier. Scale bar=50 µm. **B)** Experimental scheme of **Panels C-D**. Streptomycin pre-treated C57BL/6 mice were infected orally with 5×10^7^ CFU of wt *S*.Tm for 3 days. One group (control from **Fig. 1B-C**) treated with the vector (PBS; black symbols) and the second group with PAD4 inhibitor (GSK484; purple symbols) intraperitoneally (I.P.). **C)** Relative ranks of the tagged *S*.Tm strain abundances in feces at day 3 p.i.. **D)** Evenness score at day 3 p.i.. Upper ends of the bars indicate the median. **Panels C-D**) Pooled from 2 independent experiments for each group: control (n= 11 mice) and PAD4 inhibition (n=5 mice). Two-tailed Mann Whitney-U tests were used to compare two groups in each panel. p≥0.05 not significant (ns).

Next, we set out to test if NETosis could be blocked in the gut lumen of our mouse model. For this purpose, we used the PAD4 inhibitor, GSK484, shown to effectively reduce the formation of NETs by mouse and human neutrophils [31, 32]. Here we asked if PAD4 inhibition could prevent the observed bottleneck of the luminal *S*. Tm population (**Fig. 1B-C**). We infected mice as described in Fig. 1 using a mixture of untagged and tagged *S*. Tm (1:1000), treated them with the PAD4 inhibitor GSK484 (**Fig. 3B**; see details in **Materials & Methods**), and compared the results to the control mice from **Fig. 1**. Inhibition of PAD4 did, however, only result in a weak non-significant trend in the size of the gut lumen bottleneck (**Fig. 3B-D**; evenness score: ∼0.67 vs 0.46, p=0.44). Moreover, the pathogen densities in the feces and systemic organs did not change (**Fig. S2A-B**). Microscopy images of the infected cecum revealed that neutrophil aggregates in the gut lumen still contained detectable cit-His3+ DNA also in mice treated with the inhibitor (**Fig. 3A**). It should be noted that strategies to pharmacologically prevent NETosis are still under investigation [33], and it seems clear that our approach incompletely prevented neutrophil NETosis in the gut lumen of our *in vivo*-infection model. Nevertheless, our results show that intraluminal neutrophils are closely interacting with *Salmonella* during the acute infection, and that these neutrophils are positive for the NETosis marker cit-His3. This suggests that the release of NETs into the gut lumen occurs and might contribute to defense during *S*.Tm infection.

### Intraluminal neutrophils limit continuous insults to the epithelium and prevent massive epithelial cell loss

Our approaches so far suggested a correlation between neutrophil depletion and exacerbated epithelium response to the insult. However, they remained insufficient to establish that the protection of the epithelial barrier is attributable to intraluminal neutrophils. This can partly be explained by the limitations of our experimental infection model. Wild-type (wt) *S*.Tm utilizes TTSS-1 and -2 (encoded on *Salmonella* Pathogenicity Islands (SPI)-1 and -2, respectively) to trigger gut disease [2]. In our mouse model, SPI-2 enables long-term survival of the pathogen at systemic sites and therefore causes a typhoid-like disease at later stages (which becomes very prominent after days 3-4 p.i.). At this stage of infection, other arms of the innate immune system, including neutrophils, are active in the tissue underlying the epithelial layer (e.g., the lamina propria and submucosa) [2]. Therefore, the dramatic epithelial cell shedding we observed in neutrophil-depleted mice could be related to diminished defenses and uncontrolled pathogen growth in deeper tissues. In this case, neutrophil activity in the gut tissue might be more critical for preventing epithelial damage than neutrophils in the gut lumen. To specifically study the consequences of lack of intraluminal neutrophils on epithelial responses to *S*.Tm at mature stages of infection, we infected antibiotic pretreated mice with a SPI-2 mutant (*S*.Tm *ssaV; S*. Tm^*ssaV*^). This mutant strain is still able to trigger SPI-1 -dependent gut inflammation and the enteropathy is about as pronounced as for wt *S*. Tm infections, at least during the first 2 days p.i. [2]. However, in contrast to wt *S*.Tm, the *S*.Tm^*ssaV*^ infection does not progress into a typhoid like disease at later stages [2]. With this new model, we were able to minimize a possible contribution to epithelial cell expulsion or disruptive shedding by pathogen cells residing in deeper tissues. In contrast to our previous model (Fig. 1A), we pre-treated C57BL/6 mice with ampicillin, instead of streptomycin, which ensures more robust infection kinetics in *S*. Tm^*ssaV*^ infections (unpublished data and [34]). The mice were infected with 5×10^7^ CFU of *S*. Tm^*ssaV*^ for 3 days (experimental scheme; **Fig. 4A**).

**Fig. 4.**
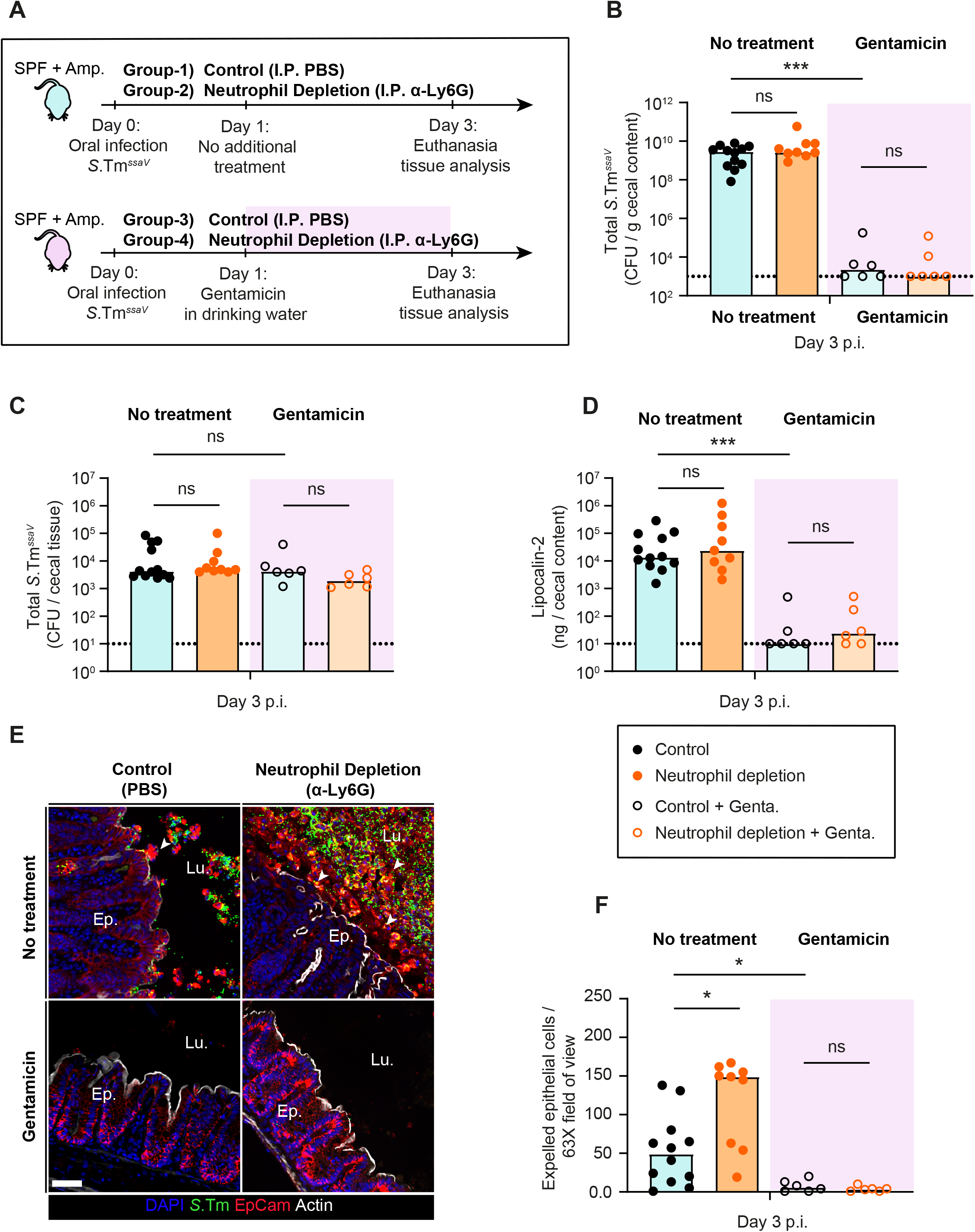
Investigation of the role of intraluminal neutrophils in a mouse model with reduced systemic disease. **A)** Experimental scheme of **Panels B-F**. Ampicillin pretreated C57BL/6 mice were infected orally with 5×10^7^ CFU of *S*. Tm^*ssaV*^ for 3 days. Mice were divided into four groups: 1) Control (I.P. PBS; black filled symbols) without gentamicin, 2) Neutrophil depletion (I.P. α-Ly6G; orange filled symbols) without gentamicin, 3) Control (I.P. PBS; black empty symbols) with gentamicin in drinking water starting from day 1 p.i., and 4) Neutrophil depletion (I.P. α-Ly6G; orange empty symbols) with gentamicin in drinking water starting from day 1 p.i. Total *S*. Tm^*ssaV*^ pathogen loads **B)** in cecal content, **C)** in the cecal tissue at day 3 p.i. in each group. **D)** Quantification of gut inflammation by Lipocalin-2 levels in cecal content. **E)** Representative micrographs of cecal tissue sections, stained for epithelial marker EpCam and *Salmonella* LPS. Lu.=Lumen. EP.=Epithelium. White arrows point at expelled epithelial cells. Scale bar=50 µm. **F**) Microscopy-based quantification of luminal IECs per 63x field of view (i.e., cells/high power field; hpf). Each data point is the average of 5 fields of view (FOV) per section. Upper ends of the bars indicate the median. **Panels B-F**) Pooled from total 4 independent experiments; at least 2 for each group: Group-1 (n=12 mice), group-2 (n=9 mice), group-3 (n=6 mice), group-4 (n=6 mice). Two-tailed Mann Whitney-U tests were used to compare two indicated groups in each panel. p≥0.05 not significant (ns), p<0.05 (*), p<0.001 (***).

To probe the significance of intraluminal neutrophils, we compared the effects of neutrophil depletion to a scenario where we removed the luminal pathogen population after day 1 p.i.. The latter was achieved by supplementing the drinking water with gentamicin, an antibiotic which acts locally in the gut lumen but does not cross the epithelium (experimental scheme; **Fig. 4A**). We reasoned that if the exacerbated epithelial response (observed in **Fig. 1D-E**) is caused by continuous high-level pathogen invasion from the luminal side, the gentamicin treatment should alleviate this outcome. First, we validated that our approach specifically removed the luminal pathogen population, while leaving tissue-lodged pathogen populations intact. At day 3 p.i., *S*. Tm^*ssaV*^ was undetectable in the cecal content in all groups undergoing gentamicin treatment while the pathogen densities stayed high in feces and cecal content throughout the experiment in groups without gentamicin (**Fig. 4B and S3A**). On the other hand, removal of the luminal pathogen population by gentamicin did not affect the pathogen loads in the cecal tissue and the systemic organs at day 3 p.i. in none of the groups infected with *S*. Tm^*ssaV*^ (**Fig. 4C and S3B**, open symbols). This provided an ideal set-up to test the role of intraluminal neutrophils. Second, we assessed if the absence of the luminal pathogen population influenced the enteric disease kinetics. Strikingly, concentrations of Lipocalin-2 (a general marker for gut inflammation) were significantly reduced at day 3 p.i. in the groups treated with gentamicin in the drinking water (both control and neutrophil depletion with gentamicin treatment; **Fig. 4D**). This suggested that the pathogen population in the gut lumen significantly contributes to the maintenance of gut inflammation. Finally, we explored the effect of neutrophil depletion on epithelial integrity. To this end, we compared the number of expelled epithelial cells in control vs neutrophil depleted mice in the absence or presence of gentamicin treatment. In the group without gentamicin treatment, neutrophil depletion caused a significant elevation of expelled epithelial cells compared to the PBS control, as we observed in Fig. 1D (**Fig. 4E-F**). Similarly, the epithelial cells again appeared to dislodge from the mucosa in an uncontrolled manner and in big aggregates in neutrophil-depleted animals (**Fig. 4E**). In stark contrast, neutrophil depletion did not lead to an elevated frequency of epithelial cell expulsion in the group receiving gentamicin treatment (**Fig. 4E-F**). In fact, the gentamicin treatment reduced the numbers of expelled epithelial cells in the lumen down to baseline at 3 days p.i. (compare open symbols to black filled symbols in **Fig. 4F**). This effect was observed in mice both with and without neutrophil depletion. Upon gentamicin treatment, the epithelium ultrastructure also recovered, as judged by clearly defined and organized crypt structures. These findings reveal that epithelial invasion events from the lumen at mature stages of *S*.Tm^*ssaV*^ infection continue to fuel epithelial cell expulsion, and that intraluminal neutrophils provide a crucial barrier to reduce the frequency of these events so that epithelial barrier integrity can be retained.

### The lack of intraluminal neutrophil defense is detrimental for the epithelium in mice lacking a resident gut microbiota

Previous studies highlighted microbiota mediated colonization resistance, mucus secretion by goblet cells, antimicrobial peptides secretion, epithelial NAIP/NLRC4 inflammasome, and neutrophil infiltration into the gut lumen as potentially complementary factors protecting the epithelium from pathogen attack. How these various protective systems are integrated with each other remains poorly understood, but it appears conceivable that defects in any of them will result in a heavier reliance on the others, e.g. a more prominent dependence on the luminal neutrophil defense predicted in animals lacking another protection factor. One good example of such a scenario would be the absence of resident microbiota, which has been shown to exacerbate *S*.Tm-induced colitis in mice [18]

To test this hypothesis, we infected germ-free mice with 5×10^7^ CFU of *S*. Tm^*ssaV*^ and compared neutrophil-depleted mice to control mice (**Fig. 5A**). Since germ-free mice lack resident microbiota, they are more susceptible to *Salmonella* infection (i.e., one of the defense layers is already missing; colonization resistance by the microbiota [35]). Importantly, the epithelial regeneration which is critical for maintaining epithelial integrity during an acute *S*.Tm infection is also delayed in germ free mice [18]. These features allowed us to scrutinize the role of intraluminal neutrophils even more readily than in antibiotic pretreated mice associated with a conventional microbiota. In germ-free mice, *S*. Tm^*ssaV*^ colonized the gut to the carrying capacity in both groups and *S*.Tm^*ssaV*^ was shed at very high numbers until day 3 p.i. (∼10^9^ CFU / g feces; **Fig. 5B**). Moreover, neutrophil depletion did not significantly affect the systemic spread of the pathogen (**Fig. 5C**). However, mRNA expression analysis of cecal tissue revealed a striking difference between depleted and non-depleted mice. Many genes encoding pro-inflammatory cytokines and chemokines associated with the acute *Salmonella* infection were induced to higher levels in the cecum of neutrophil-depleted mice compared to the controls (**Fig. 5D**). This supported our hypothesis that the absence of intraluminal neutrophils leads to an overstimulation of the mucosal immune defense.

**Fig. 5.**
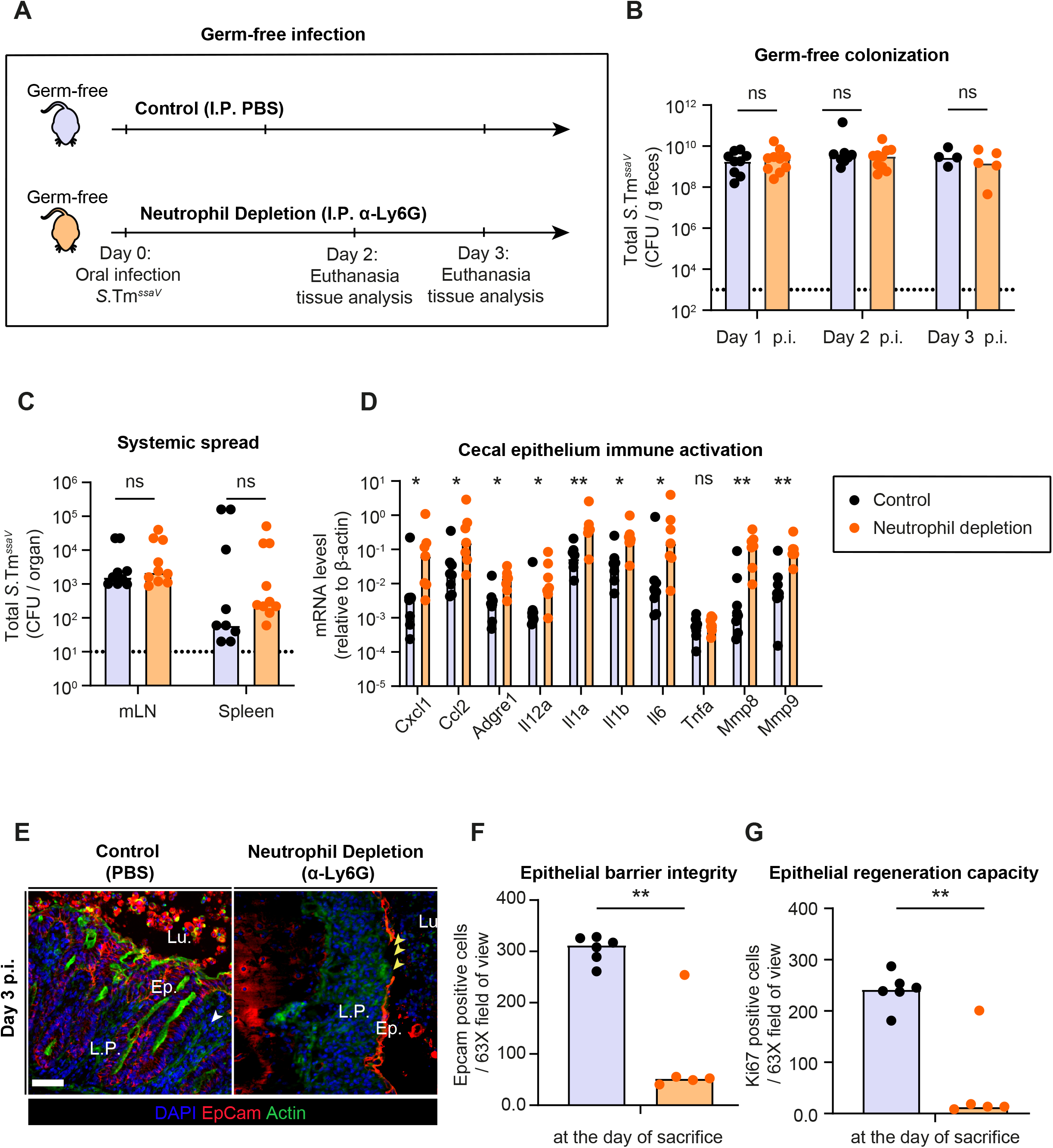
Consequences of neutrophil depletion on epithelial health during *S*. Tm^*ssaV*^ infection of germ-free mice. **A)** Experimental scheme of **Panels B-G**. C57BL/6 germ-free mice were infected orally with 5×10^7^ CFU of *S*. Tm^*ssaV*^ for 2 or 3 days (mice were euthanized at either day according to the health status of the neutrophil-depleted mice). Mice were divided into 2 groups: 1) Control (I.P. PBS; black filled symbols), 2) Neutrophil depletion (I.P. α-Ly6G; orange filled symbols). Total *S*. Tm^*ssaV*^ pathogen loads **B)** in feces, **C)** in mLN and spleen (pooled data from day 2 and 3 p.i.) in each group. **D)** Quantification of mRNA expression levels in the cecal tissue by qRT-PCR, for genes involved in immune response to acute *Salmonella* infection and genes involved in tissue remodelling. Results are represented relative to β-actin mRNA levels. **E)** Representative micrographs of cecal tissue sections, stained for epithelial marker EpCam and *Salmonella* LPS. Lu.=Lumen. EP.=Epithelium. Yellow arrows point at regions with gaps in the epithelial barrier. Scale bar=50 µm. Microscopy-based quantification of **F**) IECs per 63x field of view and **G)** Ki67+ cells per 63x field of view (i.e., cells/high power field; hpf). Each data point is the average of 5 fields of view (FOV) per section. Upper ends of the bars indicate the median. Data pooled from total 5 independent experiments. Data reported in **Panel B-G** is pooled from mice euthanized at day 2,3 and 4 p.i. as results were indistinguishable after day 2 p.i.. **Panel C**: control (n=5 from day 2 and n=4 from day 3), neutrophil depletion (n=5 from day 2 and n=5 from day 3). **Panel D**: control (n=2 from day 2, n=4 from day 3, and n=2 from day 4), neutrophil depletion (n=2 from day 2 and n=5 from day 3). **Panel F-G**: control (n=2 from day 2, n=2 from day 3, and n=2 from day 4), neutrophil depletion (n=2 from day 2 and n=3 from day 3). Two-tailed Mann Whitney-U tests were used to compare two indicated groups in each panel. p≥0.05 not significant (ns), p<0.05 (*), p<0.01 (**).

Next, we asked whether this hyper-activation of the mucosal immune responses is detrimental to the epithelial barrier. To test this, we first analysed the mRNA expression levels of matrix-metalloproteinases (Mmp-8 and -9) associated with inflammation-linked tissue remodelling [36]. Indeed, the neutrophil-depleted group showed an elevated mRNA expression of these genes (**Fig. 5D**). Second, we analysed the epithelial barrier integrity and the epithelial regeneration responses in these mice. Strikingly, fluorescence microscopy of cecal tissue at day 2 and 3 p.i. revealed that neutrophil depletion caused a severe loss of epithelial cells and formation of gaps in the barrier (**Fig. 5E-F**). The number of epithelial cells per crypt was also dramatically reduced and organized crypt structures were entirely lost in some mice (**Fig. 5E-F**). To test if this loss of epithelial barrier was associated with a reduced capacity of the epithelial cells to divide, we analysed the fraction of Ki67+ cells. Indeed, actively dividing cells were nearly absent in neutrophil-depleted groups while the active division and crypt hyperplasia was apparent in the control mice infected with *S*. Tm^*ssaV*^ (**Fig. 5G and S4A**). In control mice, more than 90% of the epithelial cells of a crypt were positive for Ki67. Therefore, the lack of gut microbiota appears to lead to an overwhelming attack on the epithelium during *S*. Tm^*ssaV*^ infection, which is counteracted by neutrophils. Taken together, these results suggest that in the presence of intraluminal neutrophils, epithelial integrity can be maintained by rapid epithelial proliferation even in germ-free mice, while the depletion of neutrophils is detrimental to the epithelial barrier integrity.

## Discussion

Since neutrophils are central players exerting diverse effector functions during the initial immune response to infections, a full understanding of neutrophil functions in the course of diverse infectious diseases is crucial. In the case of *Salmonella* gut infection, the role of neutrophils has not been fully elucidated. Here we tackled this question and carefully analysed the consequences of neutrophil depletion (with α-Ly6G) during the infection with *S*.Tm in different variants of the mouse model for *Salmonella* gut infection. Our findings revealed that intraluminal neutrophils provide a barrier in front of the epithelium during a critical stage of the infection and that neutrophil function is essential for preventing excessive epithelial cell loss, maintaining epithelial regeneration at a rate that preserves barrier integrity, and thus avoiding epithelium disruption (**Fig. S5A-B**).

In this report, we present an epithelium-protective role of neutrophils in three different variants of the mouse infection model for acute Salmonellosis (**Fig. 1**,**4**,**5**). The streptomycin pre-treated mouse model is often used to study acute *Salmonella* infection. It employs wt *S*.Tm (i.e., intact SPI-1 and SPI-2) that elicits enteropathy and is further characterized by the successive development of a typhoid-like disease, which can overwhelm Nramp-1-negative mouse strains at later time points (beyond days 5-6 p.i.). This parallel systemic infection makes it harder to study the role of immune effector mechanisms that are mounted after the early NAIP/NLRC4-dependent epithelial cell expulsion response [21]. Here, by combining ampicillin pretreatment and germ-free models with a SPI-2 mutant (which fails to grow to high levels at systemic sites), we could resolve the defenses protecting against luminal pathogen insults after day 1 p.i.. Because the mutant pathogen was defective in its ability to grow at systemic sites, we could focus on the luminal events, providing us with vital information regarding the neutrophils’ function in the gut lumen. Furthermore, this model revealed that the immune response to the pathogen is highly regulated (i.e., rapid reduction of Lipocalin-2 secretion upon antibiotic treatment; **Fig. 4D-E**). We believe that, although we use a mutant strain in a murine model, these findings may generalize to *Salmonella* gut infections in the wild, which rarely result in detectable systemic infection [37].

Our findings highlight the multi-layered nature of the mucosal defence against enteric *S*.Tm infection. In our work, the degree of epithelial damage was different depending on the type of infection model applied. These differences were particularly striking when comparing the ampicillin pre-treated mice in **Fig. 4** with germ-free mice studied in **Fig. 5**. During infection of ampicillin pre-treated conventional mice with *S*. Tm^*ssaV*^, neutrophil depletion caused excessive epithelial cell loss, but the crypts stayed intact without any larger gaps in the epithelium. In contrast, neutrophil-depleted germ-free mice developed pronounced epithelial damage and essentially a full collapse of crypt ultrastructure. This highlights the importance of resident microbiota in complementing the here described intraluminal neutrophil defense. Presumably, this is attributable to sub-acute stimulation of innate immune responses by molecular patterns derived from the resident microbiota [38]. Such sub-acute stimulation might enhance the regenerative capacity of the gut epithelium and thereby prevent pronounced epithelium disruption even if neutrophils are depleted. Therefore, the effects of antibiotic-mediated depletion of the gut microbiota should be studied in more detail to gain a deeper understanding of the role of continued sub-acute stimulation of mucosal defences by the microbiota or their loss during antibiotic treatment. Several alternative mouse models which permit *S*.Tm gut infections in the face of a pre-existing microbiota, such as low-complexity microbiota-bearing OligoMM^12^ mice or transient diet shift models, have been proposed recently [39, 40]. We hypothesize that studying the host immune responses to *Salmonella* infection in models with milder microbiota perturbation can provide us with important new insights regarding microbiota – innate immune defense integration.

Neutrophil extracellular traps (NETs) are proposed as a potent effector mechanism deployed by neutrophils to kill bacteria [26]. In the past two decades, several reports highlighted that NETs (that is host DNA decorated with antimicrobial agents) are not only a potent antimicrobial mechanism, but that NETs can also cause tissue damage and elicit autoimmunity in many diseases [27]. These reports challenged the idea that NETs are a highly specific anti-infective defense mechanism, as there are only few cases where NET release occurs without negative side-effects. Our observations suggest that the intestinal lumen may be a unique site for deploying NETs as a specific anti-infective defense, without risking NET-mediated tissue damage or autoimmunity. Our results show that aggregates of neutrophils in the gut lumen are positive for one of the NETosis markers, citrullinated histone 3. Although our approach to demonstrate a functional role of NETosis in the defense against enteric *S*.Tm infection only showed a weak trend, our results still draw an interesting image: intraluminal neutrophils may release NETs to physically trap and possibly kill enteric pathogens. Due to their accumulation on the luminal surface of the intestinal mucosa, this defence would be directed specifically against motile *S*.Tm cells [41, 42] that are in the process of swimming towards the epithelium. Moreover, it is tempting to speculate that the entrapped *S*.Tm cells could then be transported and excreted with the fecal flow. Further research is required to demonstrate if indeed NETs function as a “death-plus-disposal” trap for *Salmonella*. Nevertheless, our study does substantiate that neutrophils form large intraluminal aggregates that physically and/or biochemically shields the epithelium for sustained pathogen attack. These findings support and extend earlier work on how neutrophils can confine pathobiont expansion by intraluminal casts following a primary infection [25].

In summary, we have presented evidence for the protective role of intraluminal neutrophils during key stages of acute *Salmonella* infection in mice, by providing a protective barrier against luminal pathogen attack and hence giving the epithelium time to recover through crypt cell proliferation. Of note, our findings are limited to the murine models described in this study, and further research will be needed to assess if intraluminal neutrophils function in a similar fashion in other relevant scenarios and hosts. Our findings provide a basis for future research to disentangle the molecular mechanism(s) by which neutrophils exert this protective role, and to explore if these observations can be broadly applied across the diversity of enteroinvasive infections.

## Acknowledgements

We would like to thank members of the Hardt lab and Slack Lab for helpful discussions. We acknowledge the staff of the ETH Zürich mouse facility EPIC/RCHCI (especially Manuela Graf, Katharina Holzinger, Dennis Mollenhauer, Sven Nowok & Dominik Bacovcin) and the staff of the Microbiology Institute. Valuable comments on the manuscript by Luca Maurer and Erik Bakkeren are gratefully acknowledged. This work has been supported by grants from the Swiss National Science Foundation (310030_192567, NCCR Microbiomes), and the Monique Dornonville de la Cour Foundation to WDH.

## Author contributions

Conceptualization: EG, WDH. Methodology: EG, SAF. Investigation: EG, SAF, BDN, AH, MF, MES, WDH. Technical assistance: AH, MB, MF. Writing - Original Draft: EG. Writing - Review & Editing: EG, MES, WDH. Visualization: EG. Funding acquisition: WDH.

## Declaration of interests

The authors declare no conflicts of interest.

## Supplementary Figure captions

**Fig. S1.**
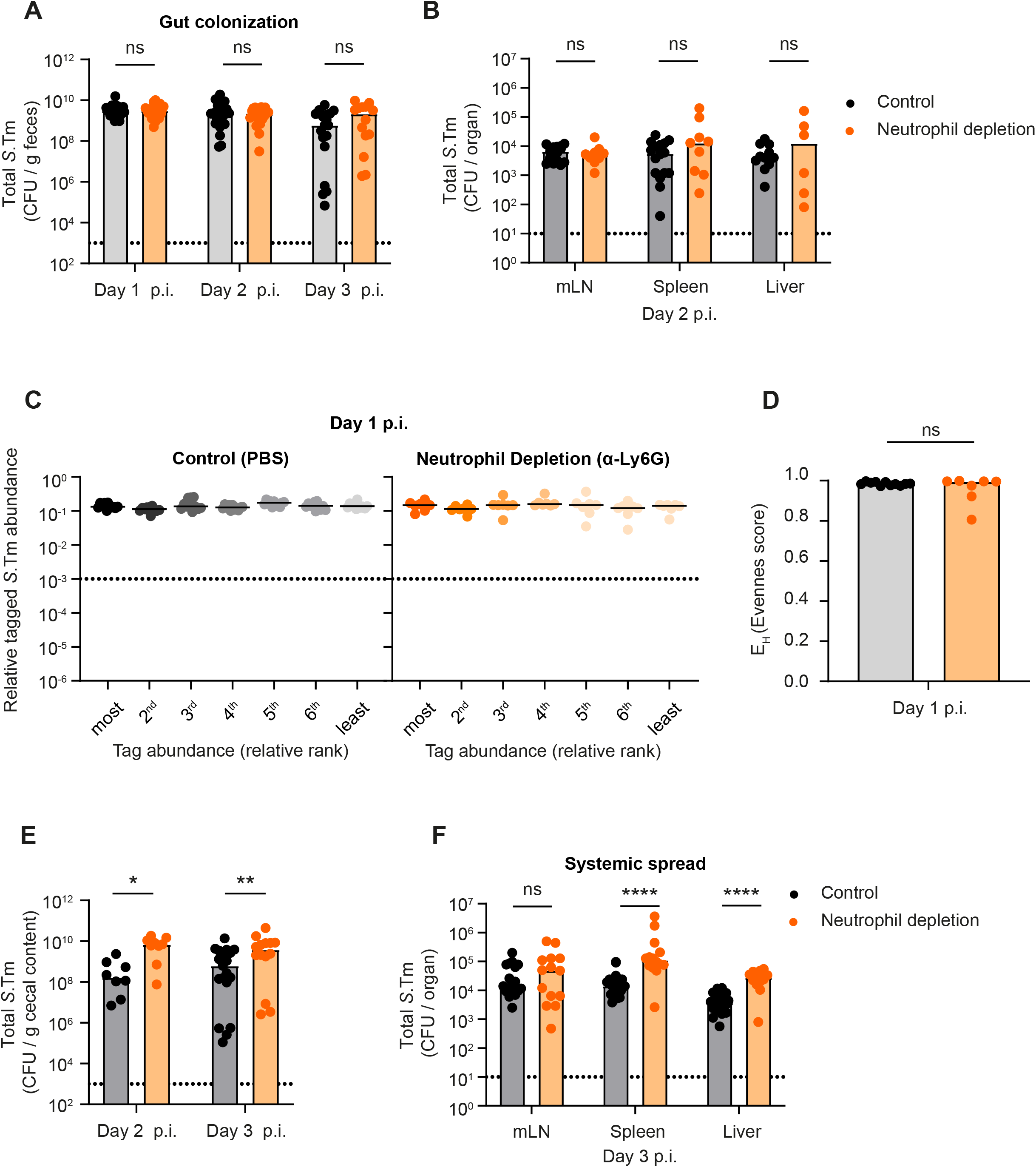
Consequences of neutrophil depletion with α -Ly6G on luminal pathogen loads, systemic spread, and epithelial cell expulsion (related to Fig. 1) **A)** Streptomycin pre-treated C57BL/6 mice were infected orally with 5×10^7^ CFU of wild-type *S*. Tm (SL1344) for 3 days. One group (control) was treated with the vector (PBS; black symbols) and the second group with α-Ly6G (orange symbols) intraperitoneally (I.P.). *S*.Tm pathogen loads **A)** in feces until day 3 p.i. (CFU / g) and **B)** in systemic organs at day 2 p.i. (CFU / organ). **C)** Relative ranks of the tagged *S*.Tm strain abundances in feces at day 1 p.i.. **D)** Evenness score at day 1 p.i. **E-F)** *S*.Tm pathogen loads **E)** in the cecal content at day 2 and 3 p.i. (CFU / g) and **F)** in systemic organs (mLN, spleen, and liver) at day 3 p.i. (CFU / organ). Lines or Upper ends of the bars indicate the median. Dotted lines indicate the detection limit. **Panel A**) Pooled from 9 independent experiments for each group: number of mice each day differs as experiments terminated at day 1, 2 or 3 p.i., but n=at least 14 at each day. **Panel B**) Pooled from 4 independent experiments for each group: number of mice for each organ differs, but n=at least 6 for each organ. **Panel C-D**) Pooled from 2 independent experiments for each group: control (n=11 mice) and neutrophil depletion (n=7 mice). **Panel E**) Pooled from at least 3 independent experiments for each group: number of mice for each day differs, but n=at least 8 for each day. **Panel** **F**) Pooled from at least 5 independent experiments for each group: number of mice for each organ differs, but n=at least 14 for each organ. Two-tailed Mann Whitney-U tests were used to compare two groups in each panel. P≥0.05 not significant (ns), p<0.05 (*), p<0.01 (**), p>0.0001 (****).

**Fig. S2.**
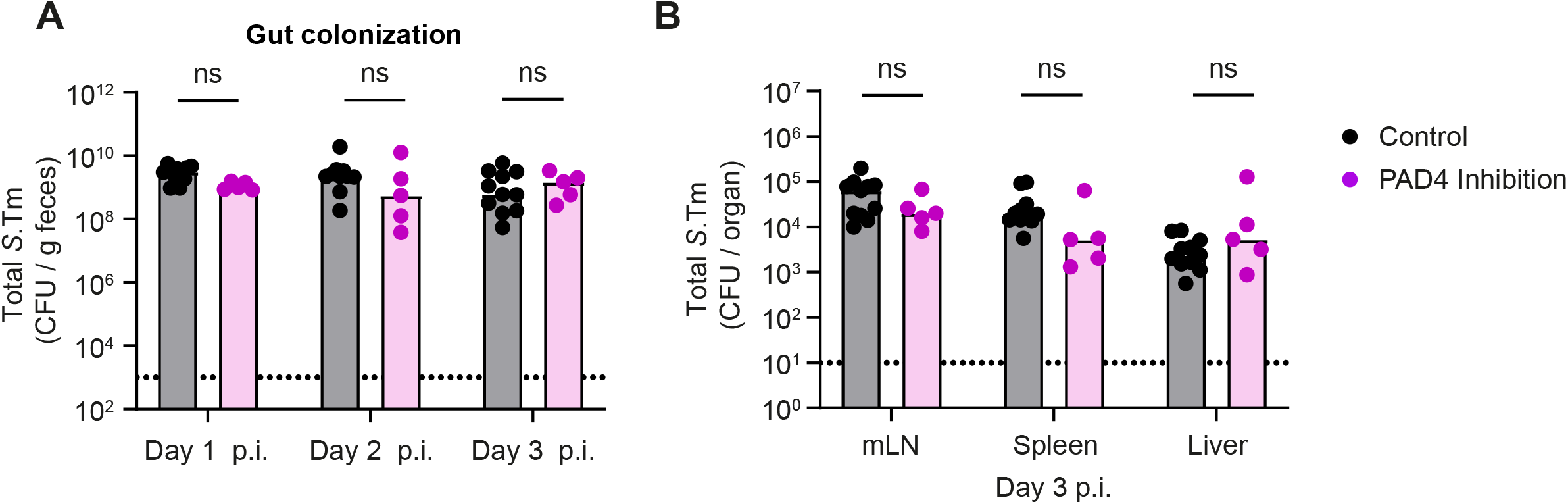
Consequences of PAD4 inhibition on luminal pathogen loads and systemic spread (related to Fig. 3) **A)** Streptomycin pre-treated C57BL/6 mice were infected orally with 5×10^7^ CFU of wild-type *S*. Tm for 3 days. One group (control from **Fig. S1A and F**) treated with the vector (PBS; black symbols) and the second group with PAD4 inhibitor (GSK484; purple symbols) intraperitoneally (I.P.). *S*.Tm pathogen loads **A)** in feces until day 3 p.i. (CFU / g) and **B)** in systemic organs at day 3 p.i. (CFU (organ). Upper ends of the bars indicate the median. **Panels A-B**) Pooled from 2 independent experiments for each group: control (n= 11 mice) and PAD4 inhibition (n=5 mice). Two-tailed Mann Whitney-U tests were used to compare two groups in each panel. p≥0.05 not significant (ns).

**Fig. S3.**
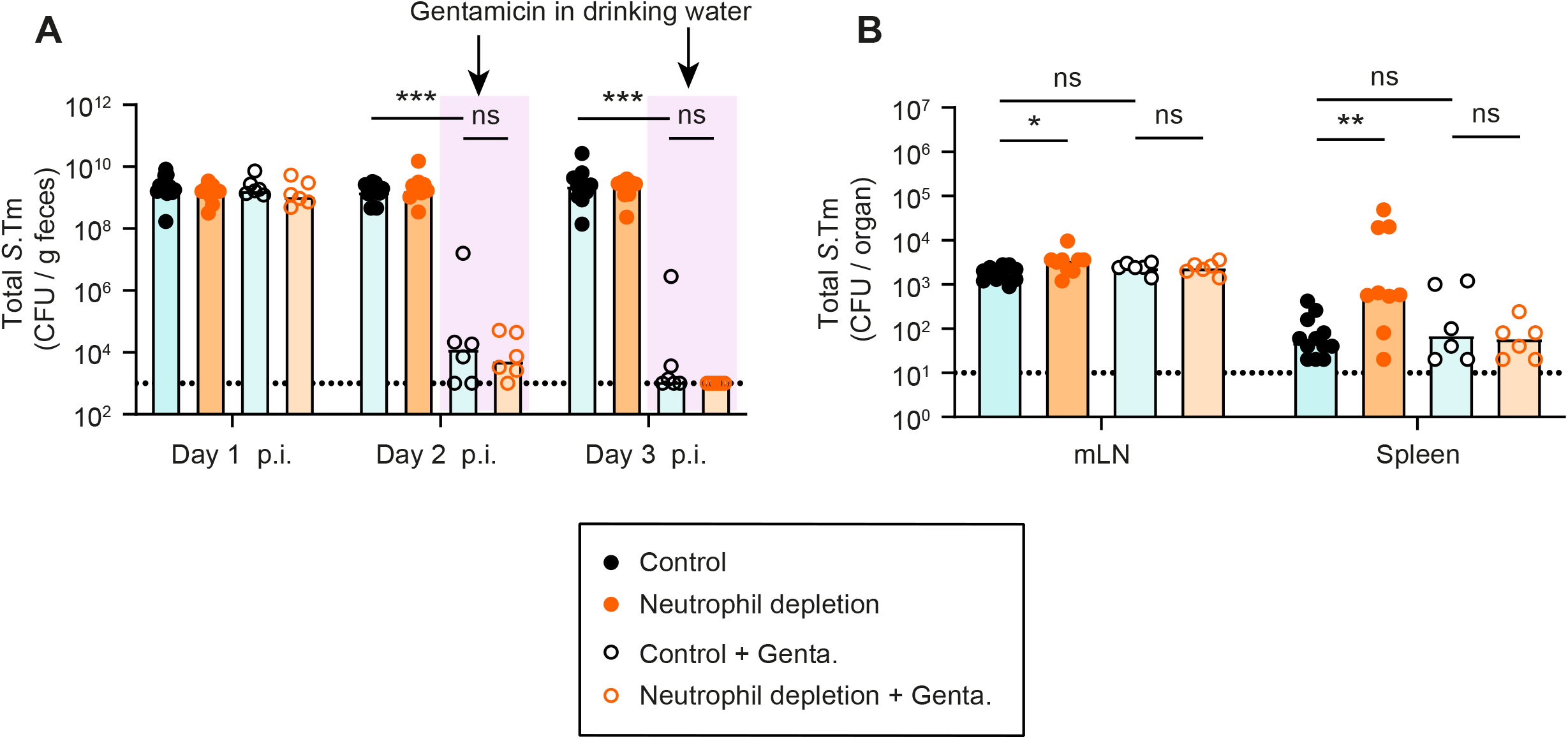
Analysis of fecal and systemic pathogen loads in a mouse model with reduced systemic disease (related to Fig. 4) Total *S*. Tm^*ssaV*^ pathogen loads **A)** in feces at day 1-3, **B)** in systemic organs (mLN and spleen) at day 3 p.i. in each group. Upper ends of the bars indicate the median. **Panels A-B**) Pooled from total 4 independent experiments; at least 2 for each group: Group-1 (n=12 mice), group-2 (n=9 mice), group-3 (n=6 mice), group-4 (n=6 mice). Two-tailed Mann Whitney-U tests were used to compare two indicated groups in each panel. p≥0.05 not significant (ns), p<0.05 (*), p<0.01 (**), p<0.001 (***).

**Fig. S4.**
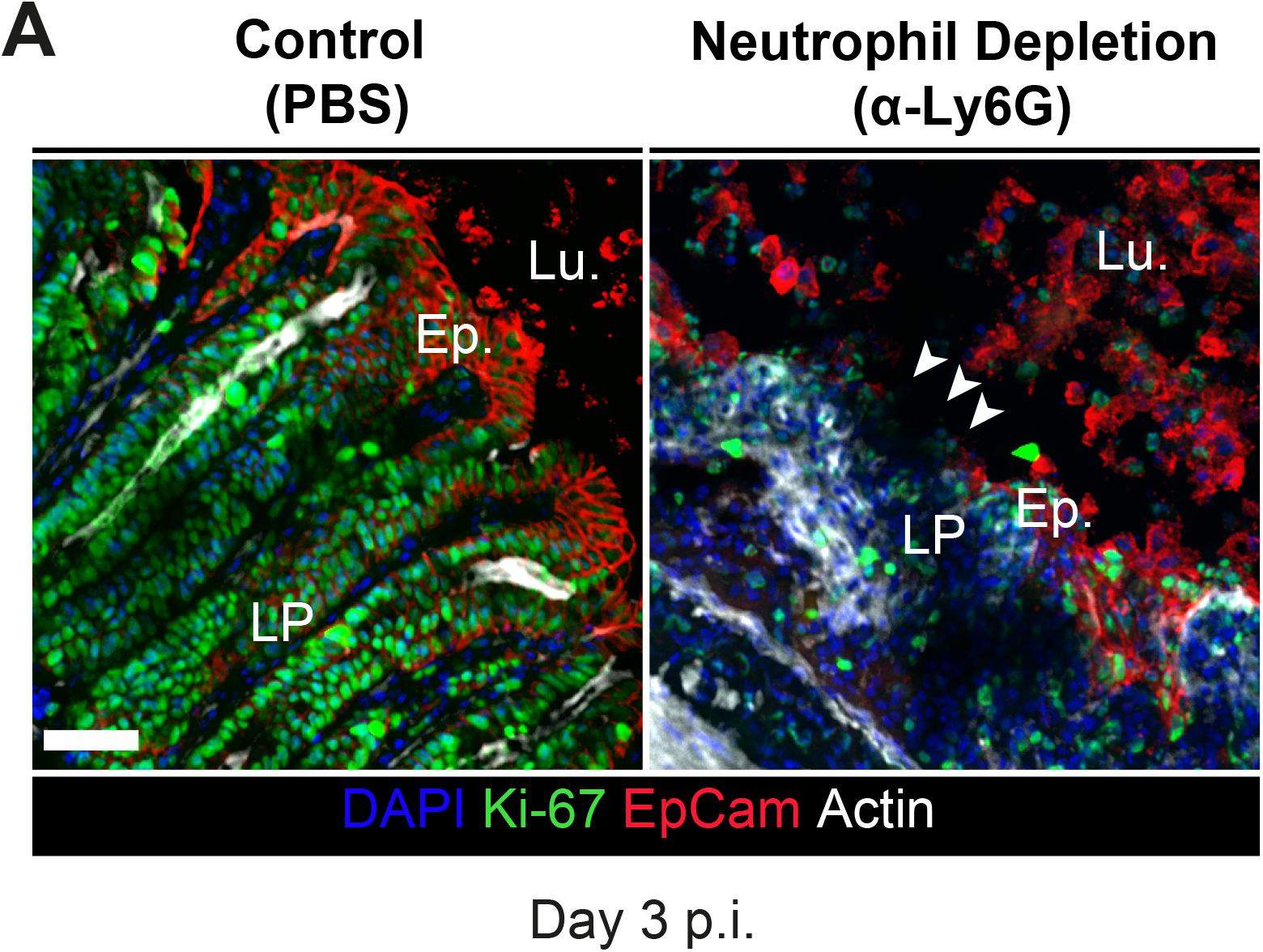
Microscopy analysis of cecal tissue in terms of its regeneration capacity (related to Fig. 5) A) Representative micrographs of cecal tissue sections, stained for epithelial marker EpCam and cell division marker Ki67. LPS. Lu.=Lumen. EP.=Epithelium. White arrows point at regions with gaps in the epithelial barrier. Scale bar=50 µm.

**Fig. S5.**
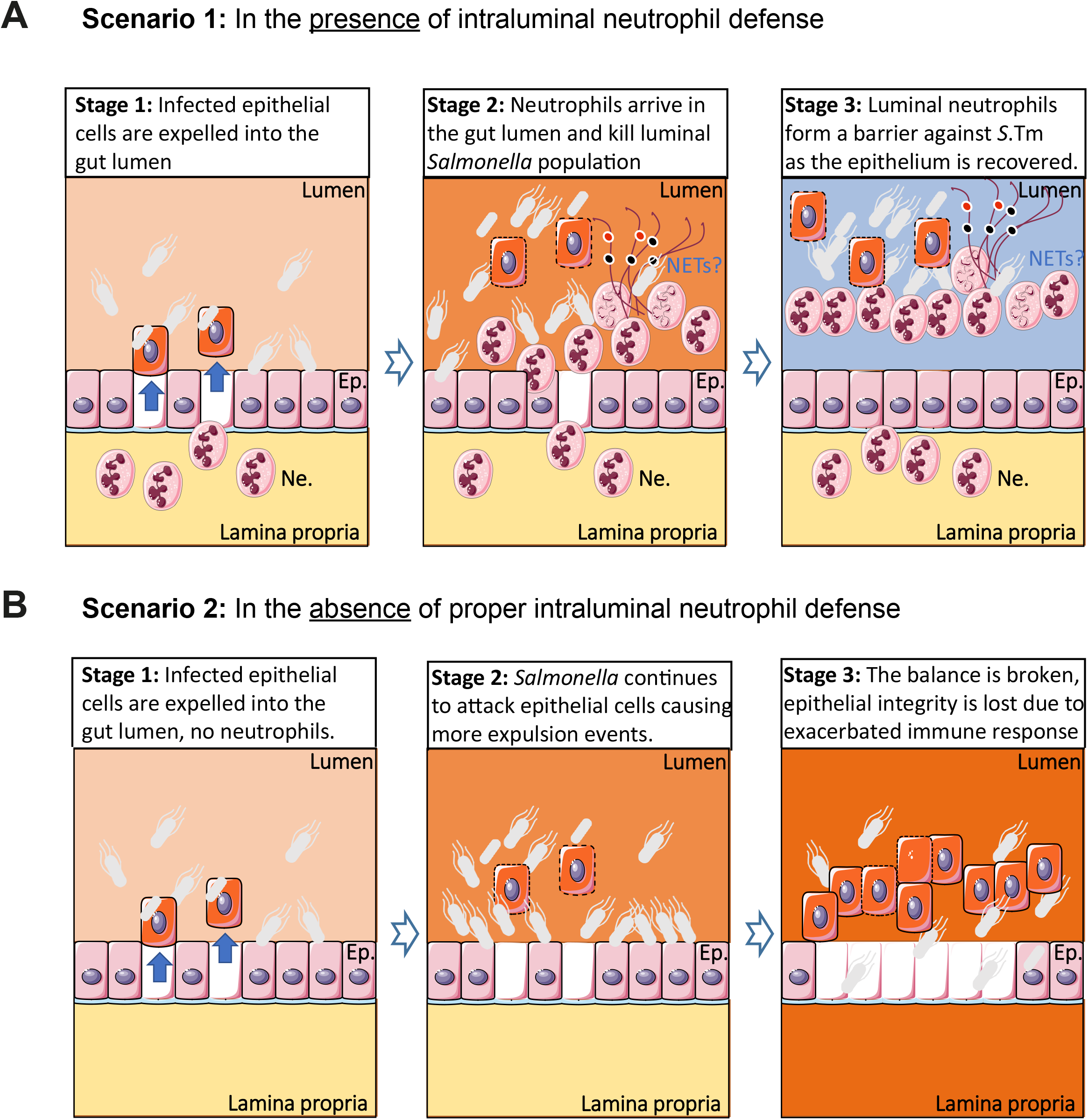
Proposed working model explaining the sequence of infection events in the presence and absence of intraluminal neutrophils. **A)** Presence of intraluminal neutrophil defense. Stage 1: *Salmonella* invasion of the epithelium is sensed by the NAIP/NLRC4 inflammasome in epithelial cells, resulting the expulsion of infected cells into the gut lumen. This leads to shortening of the crypts. At the same time, inflammasome signalling promotes recruitment of immune cells, including neutrophils, into the lamina propria. Stage 2: Neutrophils swarm into the gut lumen where they attack the invading pathogen cells and form aggregates consisting of neutrophils, NETs, and *Salmonella*. Stage 3: This barrier formed by neutrophils block further *Salmonella* attacks on the epithelium temporarily, which allows epithelial cells enough time to divide and re-establish the barrier. **B)** Absence of intraluminal neutrophil defense. Stage 1: *Salmonella* invasion of the epithelium is sensed by the NAIP/NLRC4 inflammasome in epithelial cells, resulting the expulsion of infected cells into the gut lumen. This leads to shortening of the crypts. No neutrophils are recruited. Stage 2: This leaves the epithelium exposed to further *Salmonella* attacks as the luminal population is not faced with a neutrophil counterattack. As a result, the pathogen cells continue to assault the epithelium. Epithelial cells continue to expel in response to these attacks, eventually leading to uncontrolled epithelial cell loss. Stage 3: Massive and continuous shedding result in extremely short crypts and gap formation. The epithelial barrier is breached. Underlying tissue is in direct contact with the gut luminal content.

## Materials and Methods

### Bacterial strains used in this study

In all experiments, *Salmonella* Typhimurium SL1344 (*S*.Tm;SB300; SmR) or the indicated *ssaV* mutant version (*S*.Tm^*ssaV*^; M2730; AmpR) were used. [43, 44]. WITS-tags were introduced into *S*. Tm and into by P22 phage transduction and subsequent selection on kanamycin. The presence of the correct WITS-tag was confirmed by PCR using tag-specific primers [12, 45]. For *in vivo* mouse infections, bacteria were grown in lysogeny broth (LB) media containing the appropriate antibiotics (50 µg/ml streptomycin (AppliChem); 15 µg/ml chloramphenicol (AppliChem); 50 µg/ml kanamycin (AppliChem); 100 µg/ml ampicillin (AppliChem)) at 37°C for 12h and sub-cultured in 1:20 LB without antibiotics for 4h. Cells were washed and re-suspended in cold PBS (BioConcept).

### Mouse infections

C57BL/6 mice with different microbiota complexity (germ-free and specific pathogen free (SPF)) were kept in individually ventilated cages of the ETH Zürich mouse facility (EPIC and RCHCI). Infection experiments in antibiotic pre-treated mice were done according to the previously-described Streptomycin mouse model for *S*.Tm oral infection [10]. Shortly, the mice were pre-treated with 25mg of streptomycin by oral gavage 24h prior to infection and infected on day 0 by oral gavage with an inoculum of 5×10^7^ CFU *S*.Tm. Infections in **Fig. 4** followed the same protocol but a pretreatment instead with 20mg of ampicillin and *S*.Tm^*ssaV*^ oral infection. Germ-free mice infections were done similarly but without any pretreatment. Experiments were performed with 8–12-week-old male or female mice. The sample-size was not pre-determined, and mice were randomly assigned to groups. All animal experiments were approved by the Kantonales Veterinäramt Zürich (licences 193/2016 and 158/2019).

Mice were monitored daily, and organs were harvest at the indicted time points. Feces were collected at the indicated time points and where necessary, cecal tissue and mLN were harvested at the end of the infection. For cecal tissue plating, the gentamicin protection assay was used in which the tissue is treated with gentamicin to clear extracellular bacteria. Cecal tissue was cut longitudinally, washed rapidly in PBS (3x), incubated for 45-75min in PBS/400µg/ml gentamicin Sigma-Aldrich) at RT, and washed extensively (3x 30s) in PBS before plating. For plating, the samples were homogenized with a steel ball in a tissue lyser (Qiagen) for 2 minutes at 25Hz frequency (cecal tissue 3 minutes at 30Hz). The homogenized samples were diluted in PBS, plated on MacConkey (Oxoid) plates supplemented with the relevant antibiotic(s), and placed at 37°C overnight. Colonies were counted the next day and represented as to CFU / g content. Normalized competitive index (C.I.) was calculated as the ratio of the wild type over the mutant in the feces and normalized to the initial ratio in the inoculum.

For *in vivo* depletion of neutrophils, anti-Ly6G (BioXCell, 1A8) was injected intraperitoneally at each day starting at pretreatment (In **Fig.1** 500µg/mouse, in **Fig. 4 and 5** 250 µg/mouse). For PADK4 inhibition experiments, the inhibitor GSK484 (500µg/mouse; in 10% DMSO; Cayman Chemical) was injected intraperitoneal daily starting at pretreatment.

### qRT-PCR

Cecal tissue sections were snap-frozen in RNAlater solution (Thermo Fisher Scientific) after extensive washing of the content in PBS and stored in -80°C until further analysis. Total RNA was extracted using RNeasy Mini Kit (Qiagen) according to manufacturer’s instructions and converted to cDNAs employing RT^2^ HT First Strand cDNA Kit (Qiagen). qPCR was performed with FastStart Universal SYBR Green Master reagents (Roche) and Ct values were recorded by QuantStudio 7 Flex FStepOne Plus Cycler. Primers were designed using the NCBI primer-designing tool (see **Table 1**) or ordered as validated primers from Qiagen. The mRNA expression levels were plotted as relative gene expression to β-actin (2^-ΔCt)^) and comparisons are specified in the figure caption.

**Table 1.**
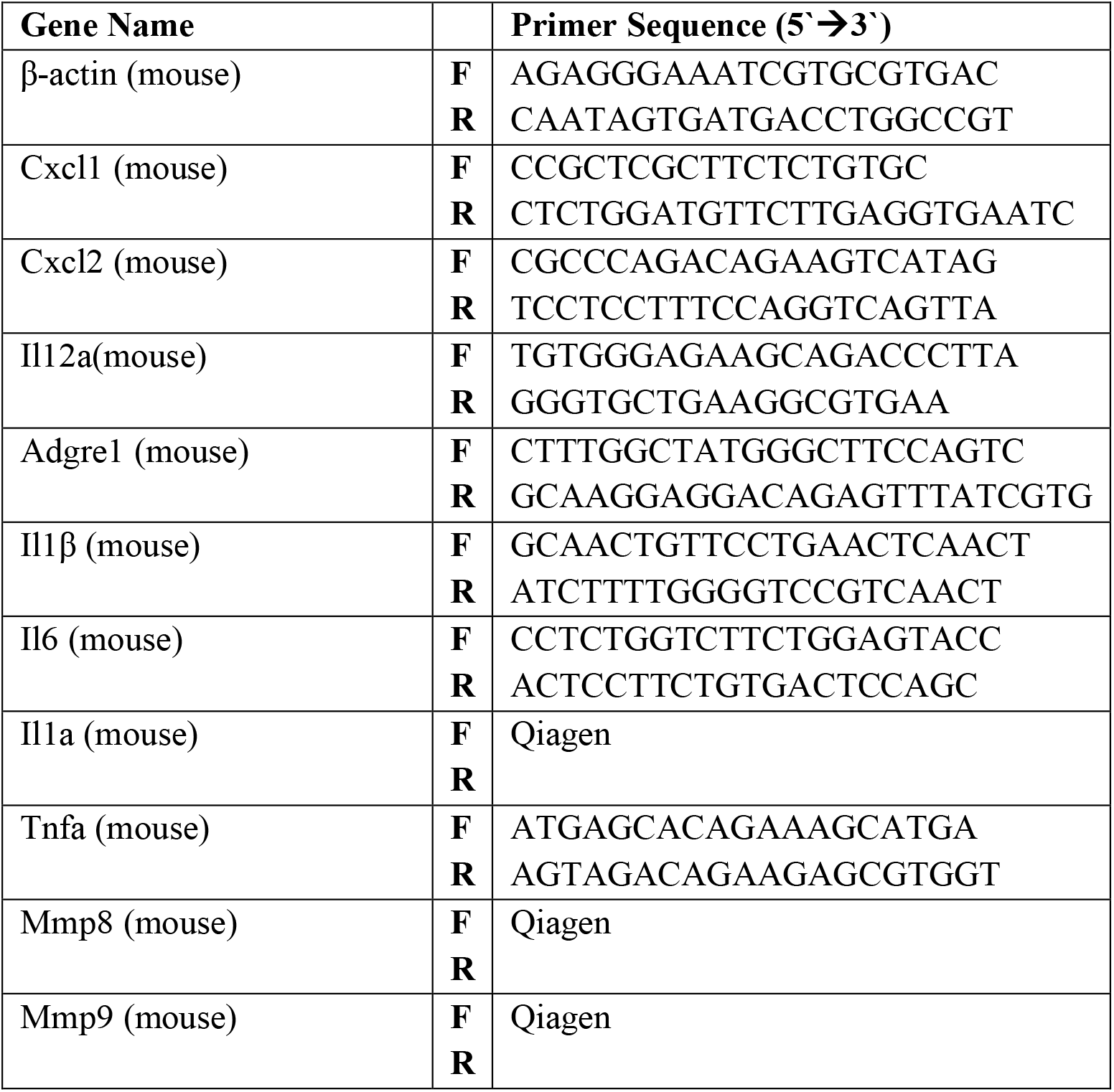
Primer Sequences used for real time qRT-PCR

### Lipocalin-2 ELISA

Lipocalin-2 ELISA (R&D Systems) was performed to determine gut inflammation from fecal samples according to the manufacturer’s instructions. Fecal pellets were suspended in sterile PBS (BioConcept), diluted 1:20, 1:400 or undiluted, and the concentrations were determined by curve fitting using Four-Parametric Logistic Regression.

### Immunofluorescence

Cecal tissue sections from mice were carefully dissected, fixed with 4% paraformaldehyde, saturated in 20% sucrose/PBS, and snap-frozen in Optimal Cutting Temperature compound (OCT, Tissue-Tek). Samples were stored in -80°C freezer until further analysis. Samples to be stained were cut in 10µm cross-sections and mounted on glass slides (Superfrost++, Thermo Scientific). For staining, cryosections on the glass slides were air-dried, rehydrated with PBS and permeabilized using a 0.5% TritonX-100/PBS solution. Sections were blocked using 10% Normal Goat Serum (NGS)/PBS before staining with primary and secondary antibodies. The following antibodies and dilutions were used for the staining of different samples: 1:200 EpCam/CD326 (clone G8.8, Biolegend), 1:200 α-*S*.Tm LPS (O-antigen group B factor 4-5, Difco), 1:200 α-Ki67 (ab15580, Abcam Biochemicals), or 1:200 α-Ly6B.2 clone 7/4, BioRad) in combination with the respective secondary antibodies, i.e α-rabbit-AlexaFluor488 (Abcam Biochemicals), α-rat-Cy3 (Jackson), fluorescent probes, i.e. CruzFluor488-conjugated Phalloidin (Santa Cruz Biotechnology), AlexaFluor647-conjugated Phalloidin (Molecular Probes), and/or DAPI (Sigma Aldrich). Stained sections were then covered with a glass slip using Mowiol (VWR International AG) and kept in dark at room temperature (RT) over night. For confocal microscopy imaging, a Zeiss Axiovert 200m microscope with 10-100x objectives or a spinning disc confocal lased unit (Visitron) with 10-100x objectives were used. Images were processed or analyzed with Visiview (Visitron) and/or ImageJ. Manual quantification was done blindly on two different sections (3 to 5 regions per section) from the same mouse according to the indicated parameters. The number of neutrophils per 63X field of view were counted on epithelium where half of the field included the lumen close to the epithelium to include freshly transmigrated neutrophils. Average crypt sizes were determined by counting the number of EpCAM positive cells per one crypt structure. Epithelial gaps were defined as the mucosal regions were where content of the lumen was in direct contact with the lamina propria. The epithelium associated *S*.Tm cells were counted by scanning the area in 5-10 µm close proximity to epithelium on the luminal side.

### Statistical analysis

Whenever applicable, the two-tailed Mann Whitney-U test was used to assess statistical significance as indicated in the figure legends. GraphPad Prism 8 for Windows was used for statistical testing. Evenness indices were calculated as previously described [12].

## Notes

### Competing Interest Statement

The authors have declared no competing interest.

